# An exploration of complex action stopping across multiple datasets: Insights into the mechanisms of action cancellation and re-programming

**DOI:** 10.1101/2024.11.03.621753

**Authors:** Sauro E. Salomoni, Simon Weber, Mark R. Hinder

## Abstract

A long history of psychological experiments has utilised stop signal paradigms to assess action inhibition. Recent studies have investigated complex stopping behaviours, such as response-selective stopping where only one component of a bimanual action requires cancellation. A current emphasis has been to utilise electromyographical (EMG) recordings to assess the temporal dynamics of action inhibition at the level of the muscle, beyond those based solely on observable behavioural events. Here, we combine EMG and behavioural data from 17 cohorts of healthy younger and older adults yielding over 42,000 response-selective stopping trials, providing unique insights into this emerging field. Expanding from past research in this area, our robust single-trial EMG analyses permit detection of cancelled (partial) and response-generating EMG bursts in both hands, revealing substantial overlaps in the distributions of timing of action cancellation and re-programming. These findings are consistent with recent experimental and modelling evidence, suggesting that response-selective stopping involves two independent processes: a discrete bimanual stop and initiation of a *new* unimanual response. This overlap appears incompatible with the recent pause-then-cancel model, and more consistent with a broader “pause-then-retune” account, where a slower process mediates any action updating, not just cancellation. Moreover, this independence means that cancellation can happen at any time during motor planning and execution, against the notion of an observable “point of no return” in motor actions. We also discuss best practices for the analysis of EMG data and indicate how methodological aspects, such as choosing appropriate reference time points, can influence the outcomes and their interpretation.

## 1. <u>Introduction</u>

Action inhibition – the ability to cancel a planned, or initiated, voluntary action – is a key component of human executive function. It is most commonly investigated using the stop signal task (SST), whereby participants respond (e.g., press a button) as quickly as possible to an imperative go signal (go trial), but need to withhold that response (stop trial) if an infrequent stop signal is unexpectedly presented after a delay (stop signal delay, SSD). According to the classic independent race model (Logan et al., 1984; Logan & Cowan, 1984), the ability to successfully inhibit or cancel the action is determined by a race between independent go and stop processes. Despite its simplicity, this model provides a suitable framework to estimate an average time of the unobservable inhibitory process (stop signal reaction time, SSRT) (Verbruggen et al., 2019). However, SSRT estimations based on the race model may not be accurate in the presence of excessive proactive slowing where the assumption of independence is violated (Verbruggen et al., 2013), and cannot account for failures to initiate the stop process (Band et al., 2003; Matzke et al., 2017). Moreover, the classic race model may also be inappropriate for complex variations in the SST that are thought to be more reflective of everyday inhibitory behaviours (Aron & Verbruggen, 2008; Coxon et al., 2007) – for example, in response-selective stopping (referred to as *selective stopping* from here onwards), where go signals typically require bimanual responses (e.g., simultaneous button presses with both the right *and* left fingers), but stop signals require cancellation of only the left *or* right response, while the other effector continues to respond.

As an alternative to SSRT estimations, analysis of electromyographical (EMG) data from task-relevant muscles enables single-trial calculations of the times of action cancellation.

Specifically, a subset of successful stop trials exhibit ‘partial’ EMG bursts (Boxtel et al., 2001; Jong et al., 1990), representing covert responses initiated following the imperative go signal, but cancelled upon presentation of the stop signal before they could generate an overt behavioural response. The latency of the peak amplitude of a partial burst relative to the time of the stop signal (SSD), termed EMG CancelTime or partial response EMG (prEMG), represents the time when EMG amplitude begins to decline and provides an *observable* (rather than *estimated* as per SSRT) measure of the time required to cancel the initial response following stop signal presentation (Jana et al., 2020; Raud & Huster, 2017). EMG CancelTime is generally reported in the range of 140-170 ms (Raud et al., 2022; Salomoni et al., 2023; Thunberg et al., 2023), broadly corresponding to the behavioural estimates of SSRT (∼200 ms) once electro-mechanical delay is taken into account – i.e., the time between the onset of muscle activation and force generation, typically ∼60 ms (Schmid et al., 2019). Importantly, estimations of stopping latency from each stop trial with an observable partial burst allows a more nuanced approach to assess inhibitory speed by yielding a distribution of single-trial values, instead of a single point SSRT estimate representing *all* stop trials. Moreover, EMG CancelTime can be considered a more direct measure of the physiological inhibitory process (at the level of the muscle), compared with SSRT that represents an indirect, averaged behavioural measure contingent upon properties of both muscles and experimental equipment (e.g., button stiffness, type of response action required, etc).

Despite the recent interest in single-trial estimations of EMG CancelTime, particularly in selective stopping tasks, a limitation of previous methods is that partial responses could only be detected in the stopping hand (i.e. hand cued to stop), but not in the responding hand, which is required to continue to respond unilaterally (e.g. Jana et al., 2020; Raud et al., 2020). In selective stopping, successful (selective) cancellation often results in cancellation of the initial bimanual response, prior to initiation of a *new* unimanual action (Gronau et al., 2024; MacDonald et al., 2014, 2017; Salomoni et al., 2023), hence assessing the relative timing of both the stopping and re- programmed responses is key for understanding the underlying neural and psychological mechanisms of complex inhibition. Accordingly, we recently developed EMG algorithms that allow independent identification of partial and response-generating bursts in both hands, while also imposing strict constraints on the timing and amplitude of partial bursts to distinguish them from spurious EMG or mirror activity. Using this methodology, we found substantial trial-to-trial variability in the speed of both action cancellation (partial burst) and re-programming (response-generating burst, see (Gronau et al., 2024)) and provided insights into the mechanisms of selective stopping, e.g., how stopping and re-programming are affected by proactive cues (Salomoni et al., 2023). Importantly, however, estimations of EMG CancelTime rely entirely on the presence of a partial EMG burst, which are only detected in a subset (∼30-50%) of successful stop trials (Raud et al., 2022; Salomoni et al., 2023). Given that stop trials are also infrequent events (occurring on 25% - 33% of all trials), and that SSD staircasing causes participants to *fail to stop* on approximately half of these, it becomes clear that the small number of trials with a partial burst during a single experimental session is often a limiting factor in allowing reliable insights into the underlying inhibitory process given the aforementioned variability (Raud et al., 2022; Salomoni et al., 2023).

Therefore, we propose that comprehensive analyses over large (combined) datasets will help produce more robust estimates and substantially improve our understanding of the processes underlying selective stopping.

To this end, in this *Exploratory Report*, we apply our recently developed EMG methodology across multiple SST datasets collected in our laboratory, yielding over 42,000 selective stop trials from hundreds of individual participants. This large combined dataset provides a rich opportunity to provide significant insights into the processes underlying complex action inhibition, and to investigate the reliability and robustness of behavioural and EMG-based measures (see Thunberg et al. Cortex 2024). We use average EMG profiles and single-trial data to investigate the temporal dynamics of selective stopping, the relative timing of action cancellation and re-programming, and the factors influencing the presence and characteristics of partial EMG bursts. Our analyses provide a framework to critique a number of key historical and contemporary theoretical concepts within the action inhibition literature, such as the point of no return and the pause-then-cancel model of action inhibition. Furthermore, we discuss key methodological considerations for EMG analysis in selective stopping for the benefit of future research and knowledge advancement.

## 2. Methods

### 2.1. Overview of data exploration

We use (qualitative) visualisations of behavioural and EMG parameters, as well as smoothed EMG profiles, across multiple datasets to enable a detailed exploration of the processes involved in selective stopping. To this end, we present (i) *average EMG profiles*, combined across datasets using a number of different reference points for synchronising to permit visual presentations of the distinct processes contributing to selective stopping; (ii) *EMG amplitude parameters* from all single trials, for example peak amplitude and the amplitude at the time of stop signal presentation, which are then assessed to evaluate whether these features may underlie successful stopping; (iii) the relative *timing of the go and stop processes*, including RT and EMG onset relative to the go/stop signal; and (iv) *the entire distribution of EMG profiles* from all stop trials to investigate the timing of action inhibition and initiation of the re-programmed action, and highlight how these processes can merge into a single inseparable burst of EMG activity. .

### 2.2. Experimental datasets

We combined behavioural and EMG data from five different studies published by our lab group, in addition to some of our unpublished data (see summary with corresponding references in Table 1). This yields more than 42,000 stop trials and 100,000 go trials, performed by more than 200 participants in 17 different experimental cohorts, comprising healthy young and older adults. All cohorts involved selective stopping variants of the SST, covering a wide range of experimental manipulations, including button presses performed by flexion or abduction of the index fingers, presentation of proactive cues (informing which hand may have to cancel its response), presence of ignore trials (i.e., salient stimuli that had to be ignored, rather than signalling a requirement to inhibit an action; known as stimulus-selective inhibition), as well as combinations of visual and/or auditory go and stop stimuli.

**Table 1:** Description of datasets included in the analyses. All experiments involved a response-selective variation of the stop signal task. Only selective (left, right) stop trials were used for the analyses, which required inhibition of one component (hand) of an initial bimanual response.

| Reference | Cohort num | Num participant | Age | Finger task | Type of stimuli: Go / Stop | Trial types during experiment |  |  |  |  |  | Number of stop trials |  |  |
| --- | --- | --- | --- | --- | --- | --- | --- | --- | --- | --- | --- | --- | --- | --- |
|  |  |  |  |  |  | Go both | Stop both | Stop left | Stop right | Ignore left | Ignore right | Stop success | Stop fail | Stop total |
| (Weber et al. 2023)<br>Cortex | 1 | 35 | 35 ± 15.0 | Abd | Visual / Visual | x |  | † | x |  |  | 1,099 | 995 | 2,094 |
|  | 2 |  |  | Abd | Visual / Visual | x |  | † | x |  | x | 988 | 1,105 | 2,093 |
| (Weber et al. 2024a)<br>Psychophysiol | 3 | 24 | 25 ± 5.5 | Abd | Visual / Visual | x |  | x | x | x | x | 590 | 599 | 1,189 |
|  | 4 |  |  | Abd | Visual / Visual | x |  | x | x |  |  | 508 | 691 | 1,199 |
|  | 5 | 27 | 71 ± 6.0 | Abd | Visual / Visual | x |  | x | x | x | x | 773 | 576 | 1,349 |
|  | 6 |  |  | Abd | Visual / Visual | x |  | x | x |  |  | 711 | 639 | 1,350 |
| (Weber et al. 2024b)<br>J Cogn Neurosci | 7 | 31 | 31 ± 10.5 | Abd | Audio / Audio | x |  | x | x |  |  | 752 | 795 | 1,547 |
|  | 8 |  |  | Abd | Visual / Visual | x |  | x | x |  |  | 791 | 757 | 1,548 |
|  | 9 |  |  | Abd | Audio / Visual | x |  | x | x |  |  | 789 | 756 | 1,545 |
|  | 10 |  |  | Abd | Visual / Audio | x |  | x | x |  |  | 799 | 750 | 1,549 |
|  | 11 |  |  | Abd | Visual / Audio-visual | x |  | x | x |  |  | 815 | 738 | 1,553 |
| (Salomoni et al. 2023)<br>Sci Rep | 12 | 37 | 24 ± 5.2 | Flex | Visual / Visual | x |  | x | x |  |  | 3,318 | 3,765 | 7,083 |
|  | 13 |  |  | Flex | Visual / Visual | x |  | x | x |  |  | 3,454 | 3,628 | 7,082 |
| (Gronau et al. 2024)<br>Cogn Psychol | 14 | 28 | 24 ± 4.6 | Flex | Visual / Visual | x | x | x | x |  |  | 1,758 | 1,040 | 2,798 |
|  | 15 |  |  | Flex | Visual / Visual | x | x | x | x |  |  | 1,616 | 1,184 | 2,800 |
| Unpublished | 16 | 28 | > 60 | Flex | Visual / Visual | x | x | x | x |  |  | 2,149 | 631 | 2,780 |
|  | 17 |  |  | Flex | Visual / Visual | x | x | x | x |  |  | 2,009 | 791 | 2,800 |
| **TOTAL** |  | **210 \*** |  |  |  |  |  |  |  |  |  | **22,919** | **19,440** | **42,359** |

Abd: Index finger abduction. Flex: Index finger flexion.
†: This cohort included only 3 stop-left trials per subject in each condition, used as catch trials; excluded from analyses.
\*: A few participants took part in more than one study; each study is treated as independent for this analysis.

In all studies, participants were seated approximately 80 cm away from a computer screen, with their forearms pronated and resting on a desk. The experimental procedures were explained verbally to participants, with instructions also presented visually on the computer screen (see Figure 1A). All studies involved a response-selective version of the stop signal task presented in PsychoPy (Peirce et al., 2019). All trials commenced with a go stimulus to which participants initiated fast-as-possible bimanual button presses (i.e., press left and right buttons simultaneously). On a minority of trials (typically 25-33%), the go signal was followed by a stop signal typically requiring cancellation of either the left, *or* right, button press (see Table 1 for all stop conditions) while continuing to execute the contralateral response as quickly as possible. The proportion of selective stop trials was smaller in some cohorts, which involved presentation of either a bimanual-stop signal (20% selective-stop, 10% bimanual-stop) or an ‘ignore’ signal (15% stop, 15% ignore) – see Table 1; only selective-stop trials were considered within the current analyses. Aiming to achieve stop success rates of approximately 50%, the delay between the go and the stop signal (stop signal delay, SSD) was adjusted (staircased) in 50 ms increments, increasing or decreasing SSD after a successful or failed stop trial, respectively (independently for left- and right-stop trials). In most cohorts, both the go and stop signals were presented visually as left- and/or right-facing arrows or coloured circles, with distinct colours representing go or stop signals. In certain cohorts, the visual stimuli were replaced by brief auditory stimuli played via a headphone to the left and/or right ear, using clearly distinct frequencies for go and stop signals, or by a combination of audio and visual stimuli (see Table 1 for specific experimental details). Behavioural events, such as response time (RT) and stop signal delay (SSD), were recorded in PsychoPy at a rate of 1,000 Hz (1 ms resolution) for most studies, except for two cohorts where RT was locked to the monitor refresh rate, being recorded at 60 Hz (16.6 ms resolution).

**Figure 1.**
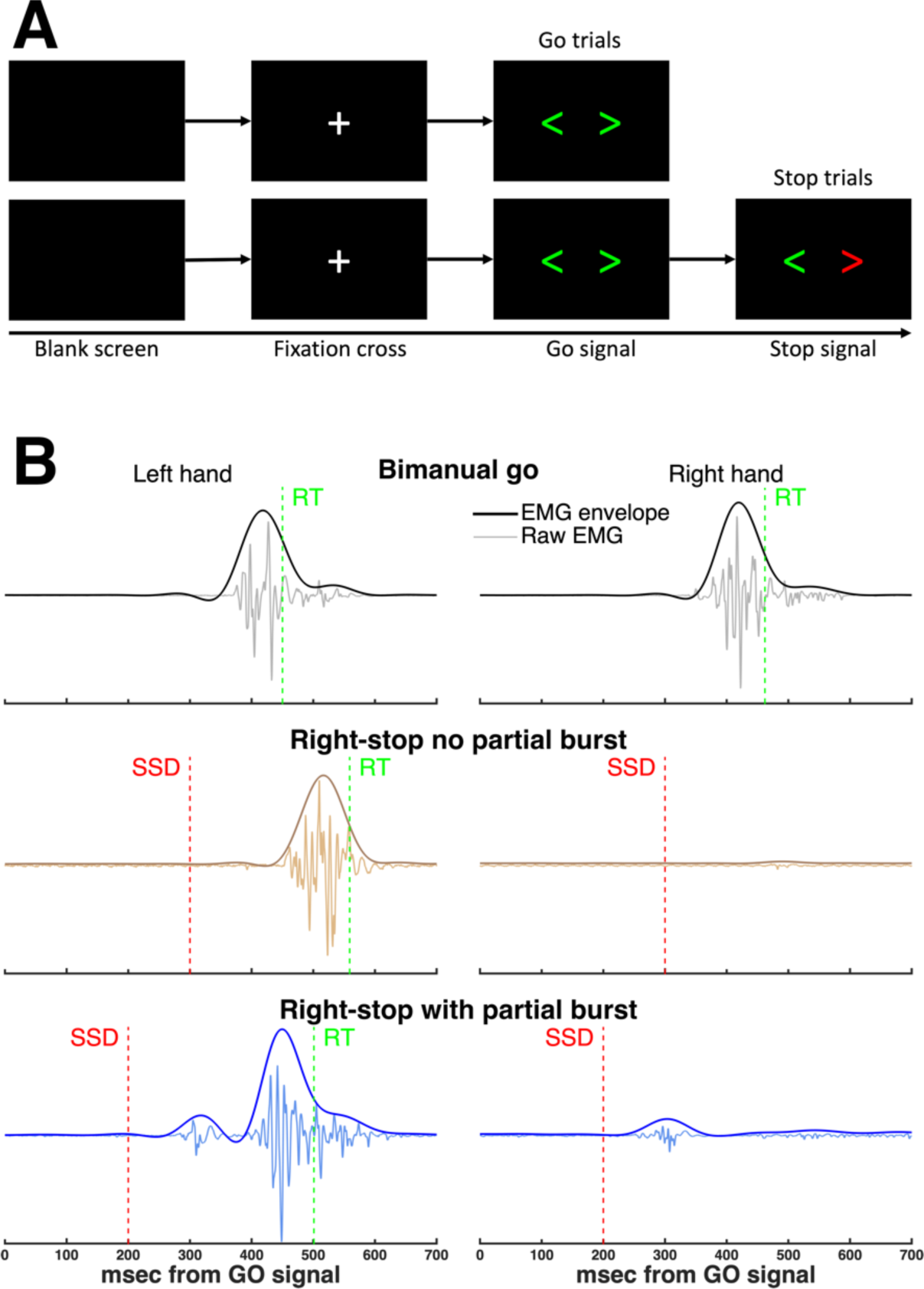
(**A**) Example of the type of visual stimuli presented during bimanual go and selective stop trials used for most of the studies included (right-stop stimulus shown in the example). Notes: In some cohorts, arrows were replaced either by circles or by a text description (“stop left” or “stop right”); in another study, visual stimuli were replaced by auditory or audio- visual stimuli. (**B**) Representative examples of raw EMG signals recorded during successful go and stop trials, and their smoothed EMG envelopes. On a proportion of stop trials, partial EMG bursts were observed (bottom row), representing cancellation of the initial bimanual response. Adapted from (Salomoni et al., 2023). RT: Response time. SSD: Stop signal delay.

### 2.3. EMG recordings

Electromyographic (EMG) signals were recorded bilaterally from the first dorsal interossei (FDI) muscles using pairs of surface Ag/AgCl electrodes (Kendall 200 series, Cardinal Health, Australia) in a belly-tendon bipolar montage, with a ground electrode on the ulnar bone of each wrist. The analogue signals were low-pass filtered at 1,000 Hz, amplified 1,000 times, recorded at 2,000 Hz (CED Power 1401 and CED 1902, Cambridge, UK) and saved to a hard drive. EMG signals were monitored throughout the experiments, and participants were instructed to relax their hands whenever tonic muscle activation was detected by the experimenter.

### 2.4. Data processing and analyses

Using behavioural data, the stop signal reaction time (SSRT) for each participant of each cohort was estimated using the integration method with replacement of omissions, averaging bimanual go-RTs from both hands, and pooling together SSDs from both left-stop and right-stop trials (Verbruggen et al., 2019). The *stopping delay*, also known as stopping interference effect (see (Gronau et al., 2024; Wadsley et al., 2022)) of each successful selective stop trial was calculated as the difference between successful stop-RT (on that trial) and the participant’s average go-RT. Any trials with RTs shorter than 100 ms were excluded from all analyses, as these likely represent anticipatory responses that were too fast to have been initiated in response to the go signal.

EMG signals were digitally band-pass filtered using a fourth-order band-pass Butterworth filter at 20 – 500 Hz. EMG profiles were obtained by full-wave rectifying and smoothing (low-pass filter) the EMG signals at 10 Hz (see Figure 1B). To allow for pooling EMG signals across participants and cohorts, single-trial EMG profiles from each participant were normalised using the peak EMG amplitude during successful go trials, averaged across all trials (separately for each hand), which also facilitates comparisons between trial types and interpretation thereof. The normalised EMG profiles were used to extract amplitude parameters, such as EMG amplitude at presentation of the stop signal (SSD) and peak EMG amplitude. EMG profiles were also used to determine EMG CancelTime, an EMG-based single-trial estimate of stopping time defined as the latency of the peak of the partial EMG burst relative to SSD (Jana et al., 2020; Raud et al., 2022; Raud & Huster, 2017), and which we have used extensively in previous studies (Healey et al., 2024; Salomoni et al., 2023; Weber et al., 2023; Weber, Salomoni, & Hinder, 2024; Weber, Salomoni, St George, et al., 2024).

As described in detail in our recent papers, we developed a robust algorithm to estimate the precise onset and offset times of the underlying muscle activity (Gronau et al., 2024; Salomoni et al., 2023) . Briefly, a single-threshold algorithm (Hodges & Bui, 1996) was used to detect all bursts of muscle activity, independently for each trial and each hand: First, the band-pass filtered EMG signals were full-wave rectified and smoothed at 50 Hz. Then, baseline EMG from each trial was determined as the segment with the lowest RMS amplitude using a moving average window with a length of 500 ms (typically the silent period before or after the button press). Finally, EMG bursts were identified as periods when the amplitude of the smoothed signal exceeded 3 SD above baseline. For robustness, any EMG bursts that were separated by less than 20 ms were merged together.

From all EMG bursts detected in each trial, we first identified the RT-generating burst, responsible for the button press, as the last burst whose onset happened (i) after the go signal and (ii) before RT. During successful stop trials, we also searched for a partial EMG burst, corresponding to bimanual responses that were initiated (after the go signal) but cancelled in response to the stop signal before generating an overt behavioural response (i.e., button press). As such, the presence and timing of these partial bursts provide an observable single-trial measure of successful inhibition of an initiated (bimanual) response (see Figure 1B, bottom row). As in our previous studies, we imposed rigorous timing and amplitude constraints for the detection of partial EMG bursts based on the theoretical assumptions of our recent SIS (*Simultaneously Inhibit and Start*) model (Gronau et al., 2024), which ensured that only partial responses related to the stop signal were included.

Specifically, our model proposes that the outcome of a selective stop trial is determined based on a ‘race’ between three distinct processes: The go signal initiates a bimanual go response; subsequent presentation of the stop signal initiates both a stop process, aimed at inhibiting the initial bimanual response, and a new unimanual go process. In successful stop trials, the fast stop process wins the race and inhibits the initial bimanual go process, leaving only the unimanual go process to win the race and register a unilateral response. Using these model assumptions, a partial EMG burst was identified as the earliest burst where (i) EMG onset happens after presentation of the go signal (i.e., initiated in response to the go signal), (ii) EMG offset happens after SSD (i.e., inhibited in response to the stop signal), (iii) peak EMG happens before the onset of the RT-generating burst from the responding hand (i.e., inhibition of the initial bimanual response precedes execution of the new unimanual response), and (iv) normalised peak EMG amplitude is larger than 10% of the reference peak EMG from successful go trials, which prevents inclusion of negligible bursts. Taken together, these constraints provide robust detection of partial responses, while preventing the inclusion of spurious EMG bursts, such as noise artifacts, mirror activity, or bursts that were initiated after RT, which are likely unrelated to the stop signal task. Note that, in two studies (cohorts 12-17) where participants pressed the buttons using finger flexion, we observed low signal-to-noise ratio (SNR) in some trials (particularly in older adults), which required the amplitude threshold (i.e., constraint ‘iv’) to be increased to 20%. In contrast, SNR was substantially increased in cohorts where buttons were pressed using finger abduction (where FDI acts as the primary agonist), thus permitting a lower threshold of 10% (cohorts 1-11).

## 3. Results and Discussion

### 3.1. Overview of experimental datasets

Figure 2A shows an overview of key timing parameters from each cohort / experimental condition, showing the entire distribution of single-trial data (or participant-level data for SSRT), as well as group averages and SD. The order of cohorts was sorted based on average go-RT to highlight its association with the other parameters. Note that the different experimental manipulations in the various cohorts (see Table 1) ensures that the distribution of each parameter covers a wide range.

**Figure 2.**
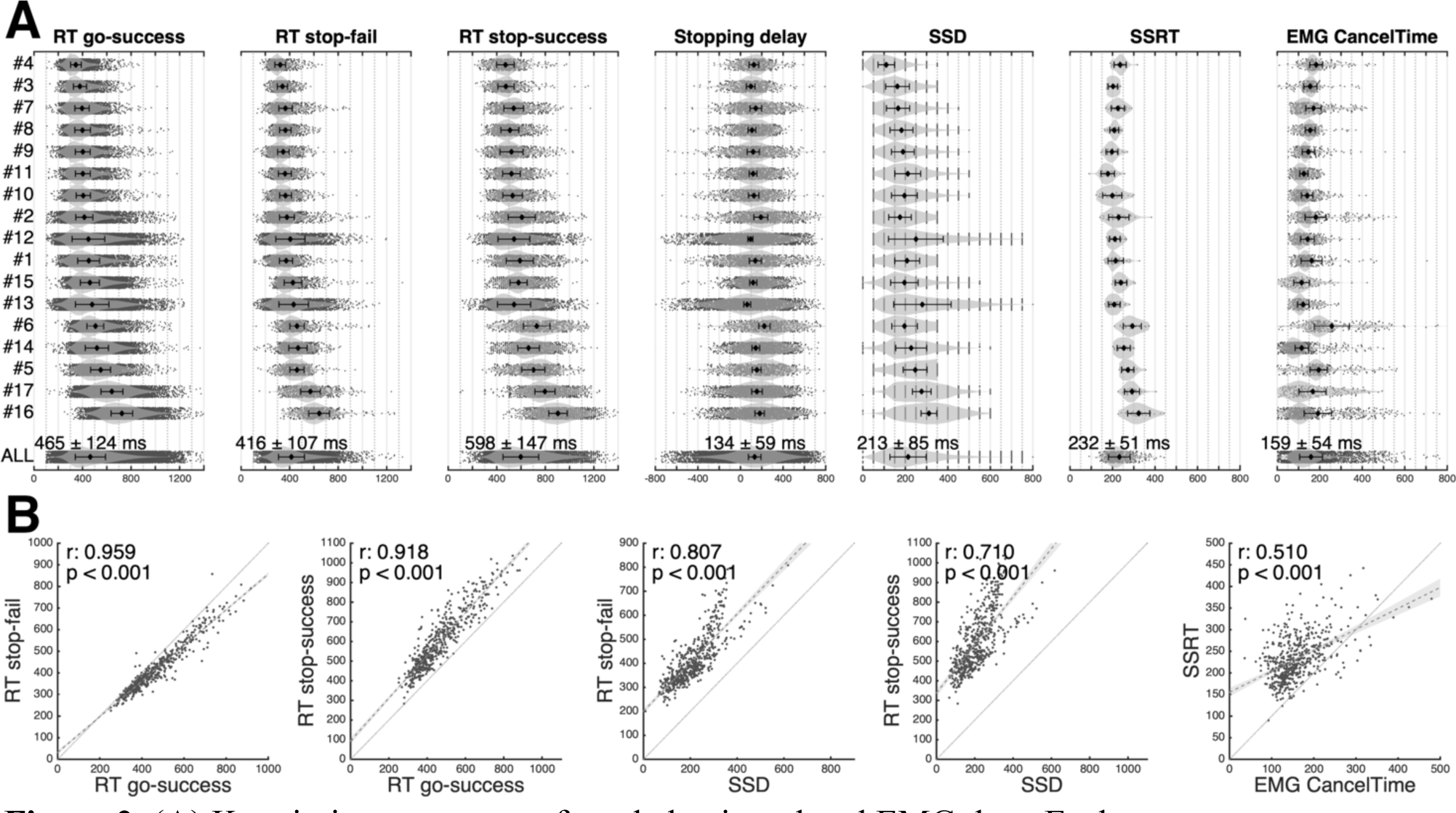
(**A**) Key timing parameters from behavioural and EMG data. Each row represents one cohort (experimental condition), sorted (top-bottom) according to their average go-RT. Numbers on the left side correspond to “Cohort num” from Table 1. Small dots represent single-trial data (or participant averages, in the case of SSRT), whereas the average from each cohort (black diamond) is shown with SD as error bars, and violin plots show the shape of the distribution, i.e., histograms. The bottom row (“ALL”) represents the conglomeration of all 17 cohorts with the grand average and SD indicated as text. Note: SSD is shown for all stop trials (successful and failed), whereas stopping delay was calculated using only successful stop trials. (**B**) Scatter plots of each participant’s average data, with regression lines (dashed; 95% CI shown as shaded area) indicating significant correlations between response times (RTs), stop signal delay (SSD), and between parameters of stopping latency, i.e., SSRT and EMG CancelTime. The identity line (dotted grey line) is shown in all plots, for reference purposes.

Visual inspection of Figure 2 reveals a typical signature of the SST: In agreement with the classic race model (Logan & Cowan, 1984), RT is shorter during *failed* stop than during successful go, as failed stops draw from the earlier part of the go-RT distribution (416 ± 107 ms < 465 ± 124 ms, r = 0.959, p < 0.001). In contrast, RT is longer in *successful* selective stop trials compared with go trials (598 ± 147 ms > 465 ± 124 ms, r = 0.918, p < 0.001), reflecting the additional time required to respond to the stop signal and execute the new unimanual response, i.e., stopping delay. Both effects can be visualised in Figure 2B: In the first column, most data points are located below the identity line, where RT stop-fail < RT go-success. Conversely, in the second column all points lie above the identity line, where RT stop-success > RT go-success. In addition, RTs from both failed and successful stop trials are strongly correlated with SSD (Figure 2B, third and fourth columns, both r > 0.7, p < 0.001). We also found a significant, albeit modest, correlation between each participant’s SSRT and mean EMG CancelTime (i.e., behavioural and EMG parameters of stopping latency, r = 0.51, p < 0.001), consistent with the associations reported in previous studies – e.g., (Raud et al., 2022; Weber, Salomoni, St George, et al., 2024).

### 3.2. Average EMG profiles: Multiple reference time points

Figure 3 shows the EMG profiles from go and stop trials, normalised to the average amplitude from successful go trials and averaged across all datasets. In each column, EMG profiles were aligned to a different reference time point: Go signal, stop signal, EMG onset, or peak EMG amplitude of the RT-generating burst. Each plot provides fine-grained details about what happens during the time near the corresponding reference time point. For example, it is clear that the normalised amplitude during the reference condition (go success) is exactly 1 when the peaks of all profiles are perfectly aligned (ref: peak EMG), but as the reference point moves to earlier time points, the peaks do not align as well, causing the shape of the EMG profiles from different trials to average out and “smooth” over time. The amount of smoothing in the average profile will reflect how well the EMG peaks align with the reference used (i.e., the distribution of relative times between peak EMG and that reference), with wider distributions resulting in greater smoothing when averaged across trials. Similarly, partial EMG bursts are clearly visible when signals are synched to the stop signal or to EMG onset, but become ‘smeared’ (and therefore less prominent) when synched to the go signal.

**Figure 3.**
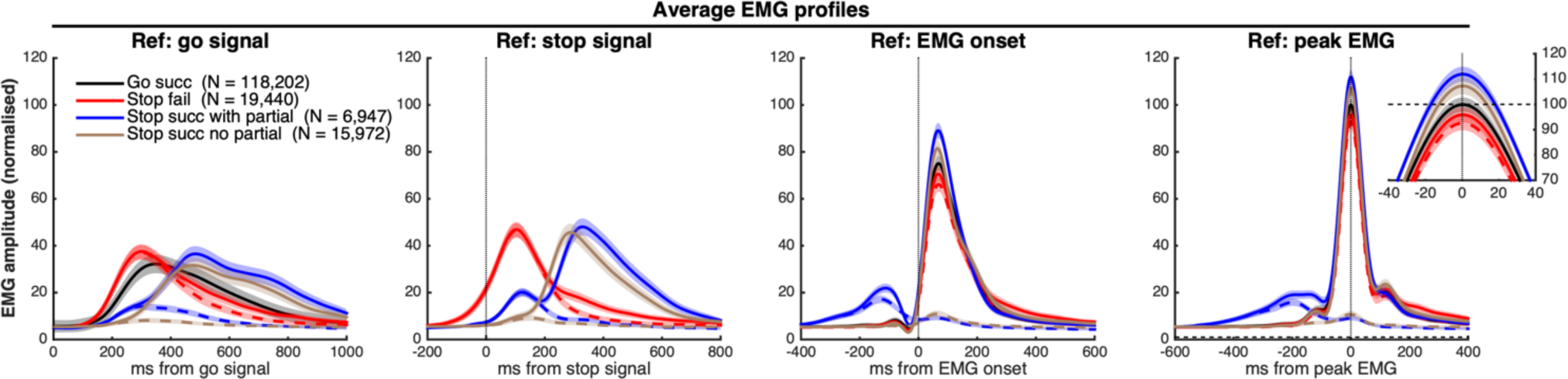
Average (± 95% CI) EMG profiles, colour-coded by trial type, stop success and presence / absence of partial bursts (see legend). Each column represents the same EMG data, but aligned to a different reference time point, resulting in different shapes of the average profiles. The stopping hand (cued to stop) is represented by dashed lines, whereas solid lines represent the responding (continuing) hand. Figure legends indicate the number of trials used to generate the average EMG profiles of each trial type. Go trials are not shown in the second column, as they do not have a stop signal.

In agreement with the behavioural data from Figure 2, when aligned to the go signal (first column), EMG responses from *failed stops* are (on average) initiated earlier than in successful go trials, whereas in *successful stops* they are initiated later. The EMG profiles aligned to the stop signal (second column) reveal that partial bursts, when present, tend to happen bimanually, as indicated by similar EMG amplitudes in both hands until ∼200 ms after presentation of the stop signal – consistent with the interpretation that they represent global (or non-specific) inhibition of the original bimanual response. Moreover, EMG amplitude at the time of presentation of the stop signal is, on average, higher during failed stops vs. successful stops with or without partial bursts. An intuitive interpretation is that stopping fails when the response is already too far underway (i.e., significant muscle activation has occurred) when the stop signal is presented, and there is not enough time to cancel the initial response before an overt response is registered.

The profiles aligned to EMG onset (third column) indicate a silent period between partial and RT-generating bursts (blue solid and dashed lines), which cannot be seen from the other reference points. Finally, plots synchronised to peak EMG (fourth column) show clear differences in EMG amplitude between trial types, highlighted in greater detail in the small inset plot. During successful stop trials, the peak EMG amplitude (of the responding hand) is, on average, ∼10% higher than during successful go trials. This observation is consistent with predictions of recent models of selective stopping (e.g., activation threshold model (ATM) (MacDonald et al., 2014, 2017), simultaneously inhibit and start (SIS) model (Gronau et al., 2024)), which predict that a stronger unimanual response is required to overcome the ongoing inhibition of the original bimanual response. If this interpretation is correct, we would expect the inhibitory input to be stronger in trials with a partial burst, where an initiated bimanual response (i.e., a partial burst) requires cancellation (Wadsley & Greenhouse, 2024); this is indeed observed, with higher EMG amplitudes during successful stop trials *with* (solid blue line) compared to *without* (solid brown line) a partial burst.

Furthermore, failed stop trials show reduced EMG amplitude in both hands, likely reflecting unsuccessful attempts to inhibit the ongoing bimanual movement. That is, failing to stop does not mean a complete absence of inhibitory input.

Taken together, these observations highlight how the choice of reference time points affects the visualisation and interpretation of EMG profiles, as the variability in relative timing strongly affects the shape of the average profiles. We recommend using multiple reference points to obtain a more comprehensive and fine-grained view of the EMG data. Indeed, without such an approach, inaccurate conclusions may be drawn in regard to EMG signatures when comparing between trial types, conditions or successful and unsuccessful stops.

### 3.3. Insights from single-trial EMG amplitudes

It has been suggested that specific EMG parameters, such as amplitude at SSD or onset time relative to SSD, can be used to distinguish between successful and failed stop trials, often discussed in the context of a *point of no return* (Fisher et al., 2024; Schultze-Kraft et al., 2016). The point of no return is a classic construct in psychology, first defined in the context of thought processes (Barlett, 1958). Since then, it has been widely used within the motor control literature to represent the point after which a planned or initiated action can no longer be cancelled, effectively dividing movements between a (controlled) preparation stage and a (ballistic or unstoppable) motor stage. Although intuitive to understand, a formal definition of the point of no return has never been established, leading to a number of disparate operationalised definitions, such as an instant in time, a locus in the sequence of processes involved in executive function, or a specific physiological threshold (Logan, 2015). Despite being loosely applied in a wide range of contexts, it remains unclear whether such a point truly exists, as there has been no unequivocal evidence of a distinct boundary between the ability or failure to inhibit a motor command. In fact, rather than a single cut-off point, recent computational model estimates using single-trial behavioural data suggest large overlaps between the distributions of finish times of the stop process (SSRTs) and the go process (stop-RTs) (Matzke, Dolan, et al., 2013; Matzke, Love, et al., 2013).

In line with the modelling evidence, although the *average* EMG profiles in Figure 3 suggests higher amplitude at SSD (i.e., the time when the stop signal is presented) during failed stops than successful stops, the *single-trial* data in Figure 4A shows a substantial overlap in the distributions of EMG amplitudes at SSD of successful and failed stop trials. This overlap provides direct evidence against a point of no return based on EMG amplitude: If amplitude at SSD was an effective predictor of whether or not the movement can be successfully stopped, we would expect to see not only a difference between average amplitudes (Figure 3), but also a separation or gap between the *distributions* of successful and failed trials. To the contrary, the histograms (Figure 4, bottom row) suggest that, although inhibition gets increasingly more *difficult* with greater EMG amplitude at SSD, it only becomes *impossible* to stop once the button is pressed (i.e., when the normalised EMG amplitude is greater than 100%, or ∼2.5% of all failed stops), and at any point prior to this, there is still a chance of successfully stopping.

**Figure 4.**
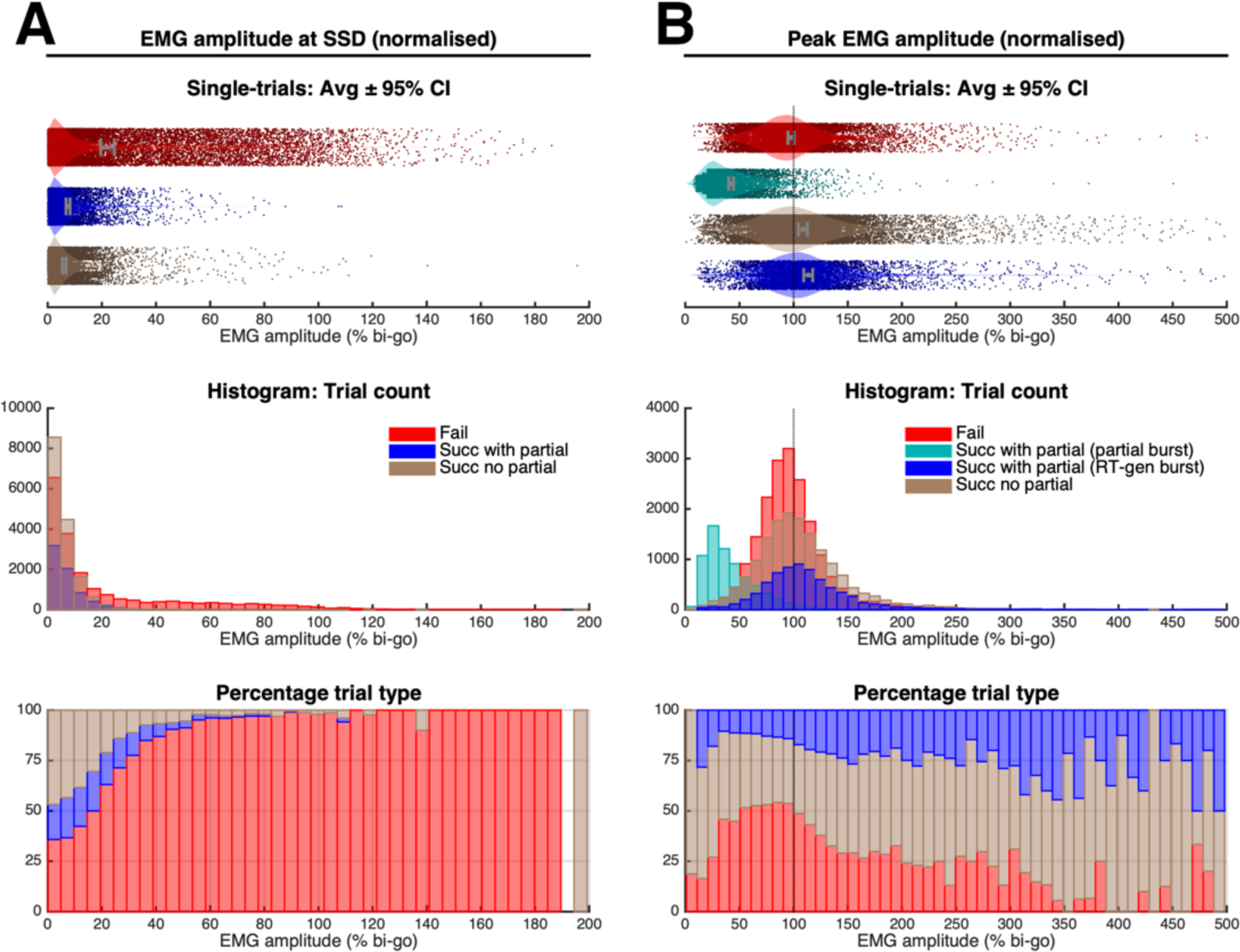
(**A**) EMG amplitude at the time of stop signal presentation (SSD); and (**B**) peak EMG amplitude. Each parameter is shown as a single-trial with average ± 95% CI (top), and histograms with trial counts (middle) and percentage of each trial type (bottom). Notes: Colour legends in the middle panels relate to data in all panels in each column. Surprisingly, a few successful stop trials (< 0.1%) had normalised EMG amplitudes at SSD greater than 100%, which likely represent either anticipated unimanual responses (i.e., “lucky guesses”), or finger movements that were misaligned or slipped off the button, precluding the registration of a button press. [If possible: 2-column figure with caption placed on the side].

The distributions of peak EMG amplitudes (Figure 4B) provide further support for the notion that unimanual responses following a successful stop are stronger than the initial bimanual response: The proportion of failed / successful stops is close to 50% for peak amplitudes between 50-100% of the bi-go amplitude, reflecting the success of SSD staircasing. However, despite the overlapping distributions, there is a steady increase in the proportion of stop trials that are successfully cancelled as peak amplitudes grow beyond 100%, suggesting that the new unimanual response requires stronger descending drive to overcome the ongoing inhibitory input associated with stopping the initial bimanual response.

### 3.4. Relative timing of the go and stop processes

In general, it is more difficult to inhibit fast responses than slow ones. In terms of response times (RTs), stopping is unsuccessful when the go response is sampled from the faster part of the distribution (Logan & Cowan, 1984), whereas successful stopping occurs when the initial go response was slower and – in the case of selective stopping – the observed RT is subject to the additional stopping delay. These effects can be clearly observed in Figure 5A, which shows the entire distribution of RTs for failed and successful stops, with and without partial bursts. On average, RTs during failed stops are shorter than during successful stop trials, as indicated by the non-overlapping confidence intervals (Figure 5A, top row). Among successful stop trials, average RTs tend to be somewhat longer when a partial EMG burst is detected (i.e., when a motor response has been initiated), presumably due to the additional time required to cancel a response that has already been activated in the peripheral muscle fibres.

**Figure 5.**
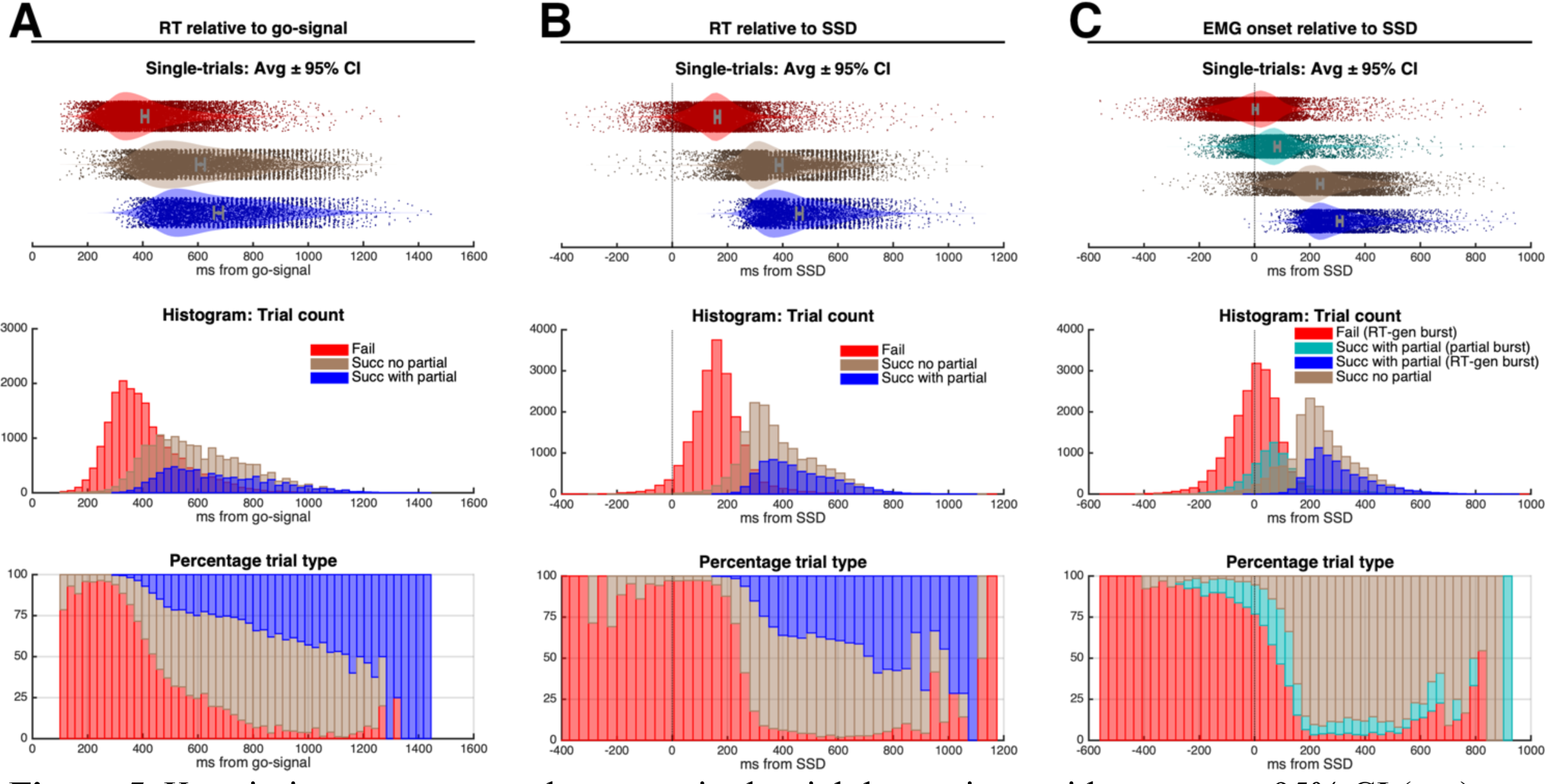
Key timing parameters, shown as single trial data points, with average ± 95% CI (top), histograms with trial counts (middle), and percentage of each trial type (bottom): (**A**) RT relative to the presentation of the go signal. (**B**) RT relative to SSD. (**C**) EMG onset relative to SSD. For failed stop trials, each data point represents the average between the left and the right hands, whereas for successful stop trials, only the responding (non-cued) hand is represented. In trials containing both partial and RT-generating bursts, trial counts of EMG onsets are shown separately for each burst, whereas only the earliest (partial burst) is shown as percentage of trial type, to avoid representing the same trial type twice. [2-column figure]

Despite the above interpretations, differences in SSD between trials and participants hinders interpretation of the RT data from Figure 5A. Accordingly, to account for individual variations in SSD, in Figure 5B we expressed RTs relative to the time of presentation of the stop signal (SSD), when the inhibitory process is initiated. This results in narrower RT distributions and a sharper distinction between failed and successful stops, while maintaining the average differences described earlier. In successful stop trials, this relative RT represents the time required to both inhibit the original bimanual response and to complete a new unimanual response – except in cases where a bimanual response was never initiated, such as (i) go signal trigger failures (Matzke et al., 2019), i.e., waiting for the stop signal before initiating a motor response; or (ii) or anticipated responses, where the participant tries to guess which unimanual response is required *before* presentation of the stop signal. Regardless, the histogram in figure 5B (bottom row) suggests that responses occurring less than 200 ms after SSD are difficult to cancel (< 5% of all stop trials are successful), which is consistent with behavioural estimates of the latency of the stop process (SSRT) being approximately 200 – 230 ms, using either the integration method (Verbruggen et al., 2019) or Bayesian modelling (Matzke, Dolan, et al., 2013; Matzke, Love, et al., 2013). That is, if action inhibition requires ∼200 ms, it is reasonable to believe that the vast majority of responses occurring within 200 ms from SSD would be impossible to stop.

Despite the insights gained by interpreting the RT distributions, behavioural responses can only provide information about the *end result* of motor activity, i.e., registration of a button press, whereas the timing of EMG onset can indicate when these actions were *initiated* at the level of the underlying muscle activity (see Figure 5C). In fact, the remarkable similarity between the distributions and timings of RT and EMG onset relative to SSD (Figures 5B and 5C), and the strong correlation between these parameters at the single trial level (stop success: r = 0.95; stop fail: r = 0.90, both p < 0.001) confirm that EMG onset times in Figure 5C represent the initiation of the same responses shown in Figure 5B, shifted by approximately 200 ms. This time shift reflects the time it takes to recruit enough motor units to generate the force necessary to press the button, in addition to the electromechanical delay. If (as suggested in Figure 5B), less than 5% of early responses (i.e., RT: [0 – 200] ms) are inhibited, we would expect to see a similarly small proportion of successful stop trials with early EMG onsets (i.e., EMG onset: [-200 – 0] ms). Instead, the histogram of EMG onsets (Figure 5C, lower panel) shows about 15-25% successful stops in this time window (Figure 5C), with most of them containing a partial EMG response, indicating that even responses initiated within 0 – 200 ms *prior to the stop signal* can still be cancelled. Indeed, Figure 5C suggests that inhibition only becomes impossible when EMG onset occurs more than 200 ms prior to SSD, likely corresponding to trials where overt responses (button presses) occurred prior to presentation of the stop signal.

### 3.5. Single-trial analyses of partial EMG bursts

We investigated the presence and timing of partial EMG bursts with respect to the subsequent RT-generating EMG bursts (responsible for overt behavioural responses). Figure 6A represents the entire distribution of EMG profiles from all successful response-selective stop trials, pooled together from all cohorts, and separated between stopping hand and responding hand. Temporal EMG profiles from each trial are represented by a horizontal line, colour-coded according to its normalised amplitude over time, and aligned to the timing of the peak of the RT-generating burst in the responding hand – similar to representations used in our previous papers (Gronau et al., 2024; Salomoni et al., 2023). To capture the timing of partial bursts (relative to the RT-generating burst), successful stops with a partial burst (in either hand) are stacked at the bottom of the plot, sorted vertically by the delay between the peaks of partial and RT-generating bursts, with a more detailed view of these EMG profiles provided in Figure 6B. Critically, these plots suggest a continuous distribution of times between the inhibited original bimanual response (partial burst) and the new unimanual response (RT-generating burst), with the estimated delay between them constrained only by the limit of our EMG onset detection algorithm (i.e., 20 ms, see Methods section 2.4). Beyond the limit of our algorithm, when only a single burst can be detected (in Figure 6A, the transition from “Successful with partial” to “Successful no partial”), a long leftward “tail” occurs in the EMG profiles at a similar latency as the latest partial bursts, suggesting a temporal overlap of the processes of bimanual inhibition and unimanual response. In an attempt to capture this “merging” between the partial and RT-generating bursts, we sorted the remaining stop trials (successful with no identifiable partial burst) based on the EMG amplitude of the “tail” prior to the peak EMG of the responding hand – more specifically, the average amplitude within the time period from -160 to -80 ms in the plots from Figure 6.

**Figure 6.**
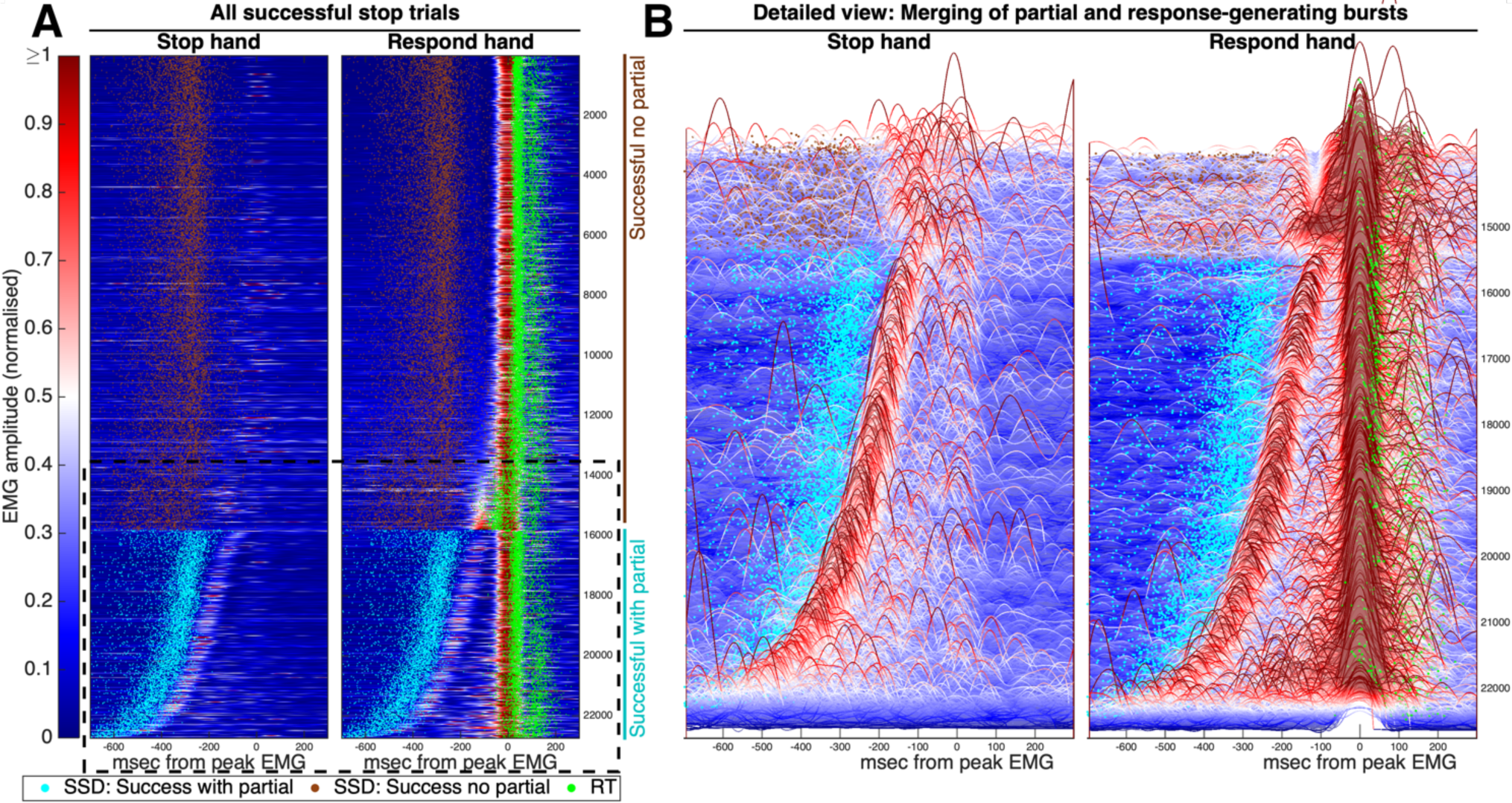
(**A**) Single-trial EMG profiles from all successful stop trials, pooled across cohorts (trial numbers shown on the y-axis). The EMG signals from each trial were colour-coded by the normalised amplitude and aligned to the peak of the RT-generating burst of the responding hand. At the bottom, trials with partial burst (in either hand) were sorted by the delay between the partial and RT-generating bursts. The remaining trials, without a partial burst, were sorted by the EMG amplitude prior to the RT-generating burst. Behavioural events (SSD, RT) in each trial are represented by small dots. The dashed black rectangle indicates the trials represented in more detail on panel B. (**B**) Detailed view of the merging between partial and RT-generating bursts. The plot is shown at a 20-degree angle from the vertical axes to enhance visualisation of the timing and amplitudes of EMG bursts from each trial. [2-column figure]

This overlap may be incompatible with the theoretical pause-then-cancel model. Originally proposed based on subcortical recordings in rodents (Schmidt & Berke, 2017), and later translated to human inhibitory control (Diesburg & Wessel, 2021), this model describes two complimentary inhibitory processes mediating action cancellation: a transient “pause” process, which is recruited broadly in response to attentional capture, and a slower “cancel” process, which can selectively remove ongoing invigoration of cortico-basal ganglia pathways. The authors interpret partial EMG bursts as reflecting the pause process, whereas the Cancel process is represented by longer-latency signatures (e.g., electroencephalographical event-related potentials occurring ∼300ms following stop signals) occurring after reduction of EMG amplitude. However, Figure 6A demonstrates that action re-programming can happen at any time following (or even during) a partial burst, which raises questions regarding the existence and/or function of a longer-latency inhibitory process. Instead, our observations support the idea of a “pause then *retune*” account (Tatz et al., 2024) where the slower process mediates action *updating* more generally (as opposed to a specific cancel process) and may either involve the initiation of a new movement, *or* further suppression of corticospinal pathways.

This observation and framework are fully consistent with our recent SIS (simultaneously inhibit and start) cognitive model (Gronau et al., 2024), which suggests the selective stop signal triggers both global cancellation and a new unimanual response. The overlap between action cancellation and re-programming observed in our data agrees with our model predictions that the distribution of finish times of the unimanual process approaches the time at which the bimanual response is cancelled. Moreover, in subsequent studies we observed similar overlaps in distributions during tasks where the “retune” process involved execution of an additional / alternative movement, rather than inhibition: (i) an “additional go task”, consisting of a unimanual choice-RT task where an infrequent visual stimulus required issuing an additional response with the contralateral hand, in addition to the original response (Weber et al., 2023); and (ii) a “change task”, where go trials consisted of a choice-RT, but an infrequent stimulus required a change of hands, i.e., stopping the initial response and responding with the other hand instead (unpublished data – manuscripts in preparation).

Furthermore, in the cohorts containing both stop and ignore trials (see table 1), we found evidence to suggest that CancelTime was similar between them (mean: stop 202 ms, ignore 204 ms, independent samples Bayesian t-test^1^: BF01 = 20.05). Because ignore cues elicit attentional capture but *do not* require action cancellation (i.e., the initiated action should still be executed), it has been argued that they may specifically involve only the pause process (Wadsley et al., 2023). However, the fact that both stops and ignore cues result in inhibition (i.e., a partial EMG burst) with the *same* latency suggests that, if this is a result of the pause process, then the pause mechanism itself also mediates the inhibition of muscle activity in response to stop trials.

The upper part of Figure 6A also shows signs of muscle activity in the stopping hand during a few (∼25%) successful stop trials in which a partial burst was *not* detected – see EMG from the stop hand around time = 0, trials labelled “Successful no partial”. The latency of this muscle activity relative to SSD (264.6 ± 173.3) was substantially longer than that observed for partial bursts (i.e., EMG CancelTime: 159 ± 54 ms), and its timing was strongly coupled with the time of the RT-generating burst from the respond hand (avg ± SD asynchrony between hands: 23.7 ± 108.1 ms).

Hence, this activity in the stop hand represents mirror activity, rather than partial responses inhibited after the stop signal. This particular observation highlights the importance of using appropriate time constraints when detecting partial EMG bursts (Gronau et al., 2024; Salomoni et al., 2023) to avoid the possibility of categorising EMG simultaneous to, or after, the response-generating burst as a partial burst indicative of action cancellation (Raud et al., 2020).

### 3.6. Detailed characteristics of partial bursts

As an extension to the analyses described above, we undertook a more detailed assessment of the prevalence, timing and amplitude of partial bursts detected during successful stop trials. The histograms in Figure 7A indicate that, when partial bursts are present, they tend to occur bimanually in at least 53% of the trials. However, it should be noted that this is likely an underestimate resulting from the stringent rules we used for the detection of partial bursts, which require normalised amplitude higher than 10%. To further investigate this, we selected all trials in which a partial burst was detected (in either hand) and looked at the EMG amplitude of both hands within a 100 ms time window centred around the time of peak partial burst. We found that the EMG amplitude (normalised to the peak EMG from successful bi-go trials) within this window was higher than 5% in *both hands simultaneously* in 89.4% of trials, which indicates the presence of bimanual EMG activity in an overwhelming majority of trials whenever a partial burst was detected in either hand.

**Figure 7.**
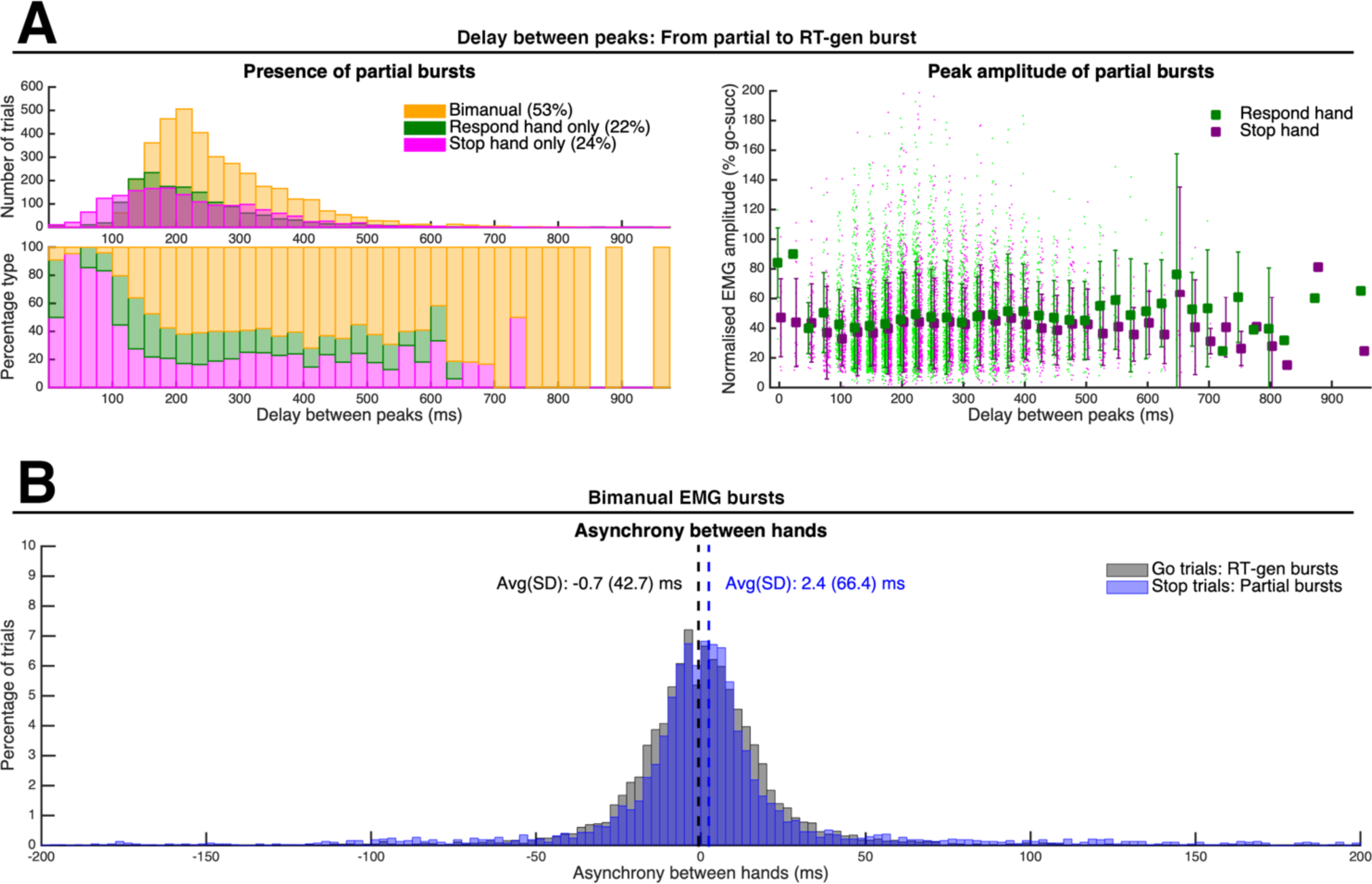
(**A**) Presence and amplitude of partial bursts as a function of the delay between the peaks of the partial and RT-generating bursts, for the responding and stopping hands. (**B**) Distribution of asynchrony between the peaks of partial bursts in the responding and stopping hands during successful stop trials, compared to the asynchrony in RT-generating bursts during successful go trials. [2-column figure]

Further evidence supporting the notion that partial bursts represent inhibited bimanual responses is presented in Figure 7B, where the difference in the timing of partial bursts (i.e., asynchrony) between hands is very small (∼2.4 ± 66.4 ms), and commensurate with the small asynchrony observed between bimanual RT-generating bursts from successful (bimanual) go trials (∼ -0.7 ± 42.7 ms).

Interestingly, as the delay between partial and RT-generating bursts gets smaller (< 100 ms), the bursts in the responding hand tend to “merge” together, and thus most partial bursts are only detected in the stopping hand (see bottom-left plot of Figure 7A). The few bursts that could be detected in the responding hand had much higher amplitudes (right column of figure 7A), consistent with the idea of merging into a single, larger burst. These larger amplitudes represent what happens at the limit precision of the EMG detection algorithm, before the partial burst becomes indistinguishable from the RT-generating component of a unilateral response.

Finally, we used correlation analyses to assess how SSD influences the characteristics of partial bursts. With longer SSDs, participants have more time to initiate their response, and the amplitude of partial bursts increase (correlation SSD vs. amplitude of partial burst: r = 0.10, p < 0.001). As a consequence, it becomes more difficult to cancel the ongoing action: Stop success rate decreases (r = -0.89, p < 0.001), and partial bursts happen less often (i.e., reduced p[partial burst | successful stop]: r = -0.89, p < 0.001), which is consistent with previous reports (Coxon 2006,Raud 2022). Moreover, in trials with longer SSDs, inhibition happens faster (i.e., reduced EMG CancelTime: r = -0.15, p < 0.001). It is possible that longer SSDs may result in less sharing of cognitive resources between the go and stop processes – however, this would violate the assumption of independence from the race model. Alternatively, shorter EMG CancelTime may indicate the presence of stronger inhibitory input. Consistent with this idea, longer SSDs were also associated with longer delays between partial and RT-generating bursts (r = 0.25, p < 0.001), i.e., longer time to re-program and initiate the new unimanual response after action cancellation. In contrast, we found no significant associations between SSD and the amplitude of the RT-generating burst (r = 0.01, p = 0.38), indicating that despite the (potentially) stronger inhibition, the strength of new response is not affected by SSD.

## 4. Conclusions and closing remarks

By combining data from multiple independent studies, this exploratory report provides a comprehensive insight into a particular aspect of complex action inhibition, namely response-selective stopping, where only part of an initial multicomponent response requires cancellation. More broadly, the qualitative approach allowed us to directly assess whether the data were consistent with a number of seminal (e.g., point of no return) and contemporary (e.g., pause-then-cancel, simultaneous inhibit and start; activation threshold) models of action inhibition. This approach yielded over 42,000 selective stop trials, providing a generalisable and robust approach, particularly when parameters with large variance, such as SSRT and EMG CancelTime, are known to be unreliable between participants and experimental sessions (Thunberg et al., 2024). The values reported here (SSRT: 232 ± 51 ms, EMG CancelTime: 159 ± 54 ms), as well as the moderate correlation between them (r = 0.49), are consistent with those reported in a recent meta-analysis that used the averages and SD of 21 cohorts from 10 independent studies (Raud et al., 2022). Moreover, the large data set allowed us to provide robust insights that could not be consistently observed across individual studies with limited trial numbers. For example, we confirmed the intuitive assumption that, in successful stops, trials with a partial burst exhibit longer RTs than trials with no partial burst, due to the additional time required to cancel a response that has resulted in (partial) activation of the effector muscles; at the individual study level, this was true in some (Gronau et al., 2024; Weber et al., 2023; Weber, Salomoni, & Hinder, 2024), but not all (Salomoni et al., 2023) of our previous studies.

Our results support the notion that, in response-selective stopping, successful response inhibition is achieved by a combination of (i) global inhibition to cancel the initial bimanual response and (ii) initiation of a new unimanual response, consistent with the assumptions of our recent SIS (Simultaneous Inhibit and Start) Bayesian computational model (Gronau et al., 2024) – see also (MacDonald et al., 2017) for the Activation Threshold Model (ATM), a compatible but distinct model framework. This conclusion is supported by studies using transcranial magnetic stimulation reporting that this global inhibition causes widespread reduction of corticospinal excitability, including that associated with the muscles of both the stopping and responding hands (Coxon et al., 2006; MacDonald et al., 2014), and even task-irrelevant muscles (Badry et al., 2009; Greenhouse et al., 2011). The inhibition raises the motor activation threshold, which effectively causes the new unimanual response to be stronger and faster than the original response (Gronau et al., 2024; MacDonald et al., 2017), as indicated by higher peak EMG amplitude and faster response time (relative to SSD) during successful stop trials compared with go trials.

From a methodological point of view, our detailed EMG analyses permit partial and response-generating bursts to be distinctly identified in both the stopping *and* responding hands. This allowed us to extend previous work (e.g., Raud et al., 2020) and conclude that the vast majority of partial bursts occur in synchrony in both hands (i.e., bimanually), consistent with the assumption that they represent the cancellation of the original bimanual response. Critically, the distribution of timing of the partial and response-generating bursts revealed a temporal overlap, where the timing of action inhibition (peak of partial burst) “merges” with the initiation of the re-programmed response (onset of RT-generating burst). This overlap suggests that the ongoing muscle activity can be modified and cancelled at any time during motor preparation or execution. Even though inhibition becomes more difficult with longer SSDs, we found that motor responses continue to be subject to voluntary control regardless of the timing or amplitude of the underlying muscle activity, and action cancellation only becomes impossible once a behavioural response is registered – against the idea of a point of no return (Jong et al., 1990; McGarry et al., 2000; McGarry & Franks, 1997) . In line with this, recent evidence accumulation models have shown improved accuracy when incorporating two distinct thresholds, corresponding to the motor (EMG) and behavioural components of the response (Servant et al., 2021).

Although EMG CancelTime provides an observable single-trial measure of stopping latency, an important limitation is that it relies on the detection of partial EMG bursts, which can only be detected in a subset (ranging from 20-70%) of successful stop trials. This effect of trial censoring means that the distribution of EMG CancelTime excludes trials where actions were cancelled before muscle activation could be detected (“cortical stop”), when participants tried to predict the upcoming trial type (“lucky guess”), or trials where the go process was not initiated (trigger failure). Therefore, in order to take full advantage of this methodology, it is important to adopt procedures that maximise the occurrence of partial bursts, including (i) using stiff buttons, which require more force (thus more EMG activation and more time) to register a response; (ii) minimising strategic slowing by regularly reminding participants to respond as fast as possible and/or providing feedback to reward fast responses; (iii) positioning the buttons sideways and asking participants to press them using (horizontal) abduction of the index fingers, which is the primary function of the FDI muscle and thus maximise the EMG signal-to-noise ratio — note that similar recommendations have also been advocated by other researchers (e.g., Raud et al., 2022). Furthermore, we also suggest imposing constraints on the timing and amplitude of partial bursts based on reasonable assumptions (for details, see Methods section 2.4) to ensure that only “real” partial bursts are included in the analysis, i.e., only partial responses that were initiated but inhibited upon presentation of the stop signal, but excluding bursts corresponding to mirror activity or other artifacts. On that note, plotting the entire distribution of partial bursts (as in Figure 6) also provides a way to visually evaluate the partial bursts detected. Our hope is that the robust methods of EMG recording and data analyses as outlined in this report will aid in our understanding of the underlying muscle activity during cancellation of complex actions.

## Credit author statement

**Sauro E. Salomoni**: Conceptualization, data curation, formal analysis, methodology, visualization, writing original draft.

**Simon Weber**: Conceptualization, methodology, writing – review and editing.

**Mark R. Hinder**: Conceptualization, formal analysis, funding acquisition, methodology, project administration, supervision, writing – review and editing.

## Data availability

All data and replication code can be made available upon reasonable request to the corresponding author.

## Footnotes

1 This independent samples t-test used a Cauchy prior distribution with a scale factor of 0.707 (Wagenmakers et al., 2018).

